# Functional investigation of a putative calcium-binding site involved in the inhibition of inositol 1,4,5-trisphosphate receptor activity

**DOI:** 10.1101/2024.08.16.608318

**Authors:** Vikas Arige, Larry E. Wagner, Sundeep Malik, Mariah R. Baker, Guizhen Fan, Irina I. Serysheva, David I. Yule

## Abstract

A wide variety of factors influence inositol 1,4,5-trisphosphate (IP_3_) receptor (IP_3_R) activity resulting in modulation of intracellular Ca^2+^ release. This regulation is thought to define the spatio-temporal patterns of Ca^2+^ signals necessary for the appropriate activation of downstream effectors. The binding of both IP_3_ and Ca^2+^ are obligatory for IP_3_R channel opening, however, Ca^2+^ regulates IP_3_R activity in a biphasic manner. Mutational studies have revealed that Ca^2+^ binding to a high-affinity pocket formed by the ARM3 domain and linker domain promotes IP_3_R channel opening without altering the Ca^2+^ dependency for channel inactivation. These data suggest a distinct low-affinity Ca^2+^ binding site is responsible for the reduction in IP_3_R activity at higher [Ca^2+^]. We determined the consequences of mutating a cluster of acidic residues in the ARM2 and central linker domain reported to coordinate Ca^2+^ in cryo-EM structures of the IP_3_R type 3. This site is termed the “CD Ca^2+^ binding site” and is well-conserved in all IP_3_R sub-types. We show that the CD site Ca^2+^ binding mutants where the negatively charged glutamic acid residues are mutated to alanine exhibited enhanced sensitivity to IP_3_-generating agonists. Ca^2+^ binding mutants displayed spontaneous elemental Ca^2+^ events (Ca^2+^ puffs) and the number of IP_3_-induced Ca^2+^ puffs was significantly augmented in cells stably expressing Ca^2+^ binding site mutants. When measured with “on-nucleus” patch clamp, the inhibitory effect of high [Ca^2+^] on single channel-open probability (P_o_) was reduced in mutant channels and this effect was dependent on [ATP]. These results indicate that Ca^2+^ binding to the putative CD Ca^2+^ inhibitory site facilitates the reduction in IP_3_R channel activation when cytosolic [ATP] is reduced and suggest that at higher [ATP], additional Ca^2+^ binding motifs may contribute to the biphasic regulation of IP_3_-induced Ca^2+^ release.

## Introduction

Changes in intracellular Ca^2+^ concentration ([Ca^2+^]_i_) control a vast array of fundamental physiological processes (1). Ca^2+^ channels, transporters, and pumps, along with Ca^2+^ binding proteins, collectively termed the “Ca^2+^ signaling toolkit”, coordinate to maintain [Ca^2+^]_i_ at low nanomolar levels under basal conditions and subsequently in response to extracellular cues to increase [Ca^2+^]_i_ with precise spatio-temporal characteristics. The localization and timing of these Ca^2+^ signals are pivotal for the appropriate activation of specific functional responses, including exocrine secretion, muscle contraction, gene transcription and cell fate decisions (2, 3). Inositol 1,4,5-trisphosphate (IP_3_) receptor (IP_3_R), are intracellular Ca^2+^ release channels and represent a critical component of the Ca^2+^ signaling toolkit. These proteins function as a major signaling hub to integrate diverse input and also facilitate the communication of Ca^2+^ signals between distinct organelles (4). The family is comprised of three distinct subtypes: IP_3_R1, IP_3_R2, and IP_3_R3 which are encoded by three different genes (*ITPR1/2/3*). These subtypes share about 60 to 70% amino acid sequence identity (5, 6). IP_3_Rs assemble as either homo- or hetero-tetramers and are predominantly located on the endoplasmic reticulum, exhibiting varied tissue-specific distribution (7, 8).

In addition to IP_3_ itself, Ca^2+^ is an obligate co-agonist of IP_3_R and is essential for channel activity (2, 9). In the presence of IP_3_, an initial increase in [Ca^2+^] facilitates IP_3_R channel opening but as the [Ca^2+^] continues to rise, IP_3_R channels close, resulting in the characteristic biphasic “bell-shaped” regulation of IP_3_R activity (10, 11). This property of IP_3_R is thought to be important for the generation of Ca^2+^ oscillations at a constant [IP_3_], known as type-1 oscillations. The initial sharp rise in the Ca^2+^ transient is due to the Ca^2+^-dependent increase in IP_3_R activity. At the peak of the transient, the elevated [Ca^2+^] reduces IP_3_R activity, allowing Ca^2+^ clearance from the cytoplasm, which alleviates IP_3_R inhibition and facilitates another cycle of IP_3_R activity. Ca^2+^ regulation of IP_3_R also contributes to the spatial characteristics of Ca^2+^ signals, including Ca^2+^ blips, Ca^2+^ puffs, and global Ca^2+^ waves, as the initial Ca^2+^ release from the most sensitive IP_3_R engages neighboring channels to propagate the signal (12–14). Notably, ATP augments the open probability of IP_3_R channels by allosterically enhancing their sensitivity to Ca^2+^, likely by binding to sites distinct from the glycine-rich Walker-A motifs, present in the linear structure of IP_3_Rs (9, 15–17). As purified IP_3_Rs reconstituted in lipid bilayers are also regulated by Ca^2+^ in a biphasic manner (18, 19), this likely indicates the presence of at least two intrinsic Ca^2+^ binding motifs within the IP_3_R structure with differing Ca^2+^ binding affinities (4, 20). Initial gel overlay assays documented ^45^Ca^2+^ binding to various IP_3_R fragments (21, 22), but since IP_3_Rs lack canonical Ca^2+^ binding motifs, identifying the domains responsible for Ca^2+^ regulation of IP_3_R activity had been elusive (23).

Recently, high-resolution cryo-EM structures of IP_3_R sub-types have identified multiple putative Ca^2+^-binding sites in rat IP_3_R1 (rIP_3_R1) and human IP_3_R3 (hIP_3_R3). Structures resolved in the presence of IP_3_, ATP, and activating [Ca^2+^] revealed densities corresponding to Ca^2+^ in a juxta-membrane domain site, formed by the third armadillo repeat (ARM3) and linker (LNK) domain of the channel. This site is referred to as Ca-IIIs in rIP_3_R1 (24) and is equivalent to the Ca-JD in hIP_3_R3 (25, 26). Mutation of this evolutionarily well-conserved Ca-III_S_/JD site, by neutralizing the negative charge of the glutamic acid residue, markedly increased the [Ca^2+^] required for IP_3_R channel activation (27). Interestingly, an analogous Ca^2+^ activation site is also conserved in the related RyR family, where similar charge mutations at this site abolished Ca^2+^-dependent activation of RyR1 (28, 29). Notably, mutating the Ca-IIIs site in IP_3_R1 did not alter the Ca^2+^-dependency for inactivation (27), suggesting that one or more additional low-affinity Ca^2+^-binding motifs may be responsible for facilitating Ca^2+^-dependent channel closure at high [Ca^2+^]_i_ (24, 27). Supporting this idea, in the presence of IP_3_, ATP, and inhibitory [Ca^2+^], densities corresponding to Ca^2+^ were reported bound to a cluster of negatively charged residues formed by the central linker domain (CLD) and the second armadillo repeat (ARM2) domain in hIP_3_R3. This site was termed the “CD Ca^2+^ binding site”, conserved across IP_3_R subtypes and through evolution but is absent in RyRs (25, 30).

To assess whether this motif serves as an inhibitory Ca^2+^ binding site in IP_3_Rs, we examined the functional impact of disrupting the conserved acidic, putative Ca^2+^ coordinating residues in the CD site on IP_3_R1 activity. We generated stable cells expressing wild type (WT) human IP_3_R1 and IP_3_R1 with charge-abrogating mutations at the CD site in either HEK-293 or DT40 cells, both lacking endogenous native IP_3_Rs (31, 32). Consistent with a reduction in inhibitory modulation of the channel, single-cell Ca^2+^ imaging assays showed that Ca^2+^ signaling in stable cell lines expressing the mutants were more sensitive to IP_3_-generating agonist compared to cells expressing WT IP_3_R. Moreover, total internal reflection fluorescence microscopy (TIRFM) revealed that mutant IP_3_R1-expressing cells exhibited fundamental Ca^2+^ release events under basal conditions and considerably higher activity following photolysis of caged-IP_3_, compared to WT cells. Further analysis of single-channel properties indicated that mutating the CD site had no significant effect on the Ca^2+^ dependency for activation or inhibition at saturating [ATP]. However, at lower [ATP] concentrations, the mutant channels displayed higher open probability (P_o_) at inhibitory [Ca^2+^] without a change in Ca^2+^ sensitivity for activation. Taken together, these findings suggest that the CD Ca^2+^ binding site plays a critical role in regulating IP_3_R1 channel activity but only at sub-saturating [ATP] levels. These results also imply that at a saturating [ATP], Ca^2+^ binding to site(s) distinct from the CD Ca^2+^ binding site is responsible for IP_3_R1 channel inactivation at high [Ca^2+^]_i_.

## Results

### The putative Ca^2+^ coordinating site is highly conserved in IP_3_R sub-types

A putative Ca^2+^-binding pocket located between the CLD and the second armadillo repeat (ARM2) domain in hIP_3_R3 is unique and evolutionarily well-conserved in IP_3_Rs (CD site) but not present in RyR (25, 27, 30) (Figs. 1A-D). Within the CD Ca^2+^ site, Ca^2+^ is coordinated by carboxyl groups from the negatively charged E1127/E1122 and E1130/E1125 in rIP_3_R1 and hIP_3_R3, respectively (Fig. 1C). In the current study, we systematically investigated the consequences of disrupting the negatively charged corresponding putative Ca^2+^ coordinating residues (E1128 and E1131) in the ARM2 domain constituting the CD site on hIP_3_R1 activity (Fig. 1A).

**Fig. 1.**
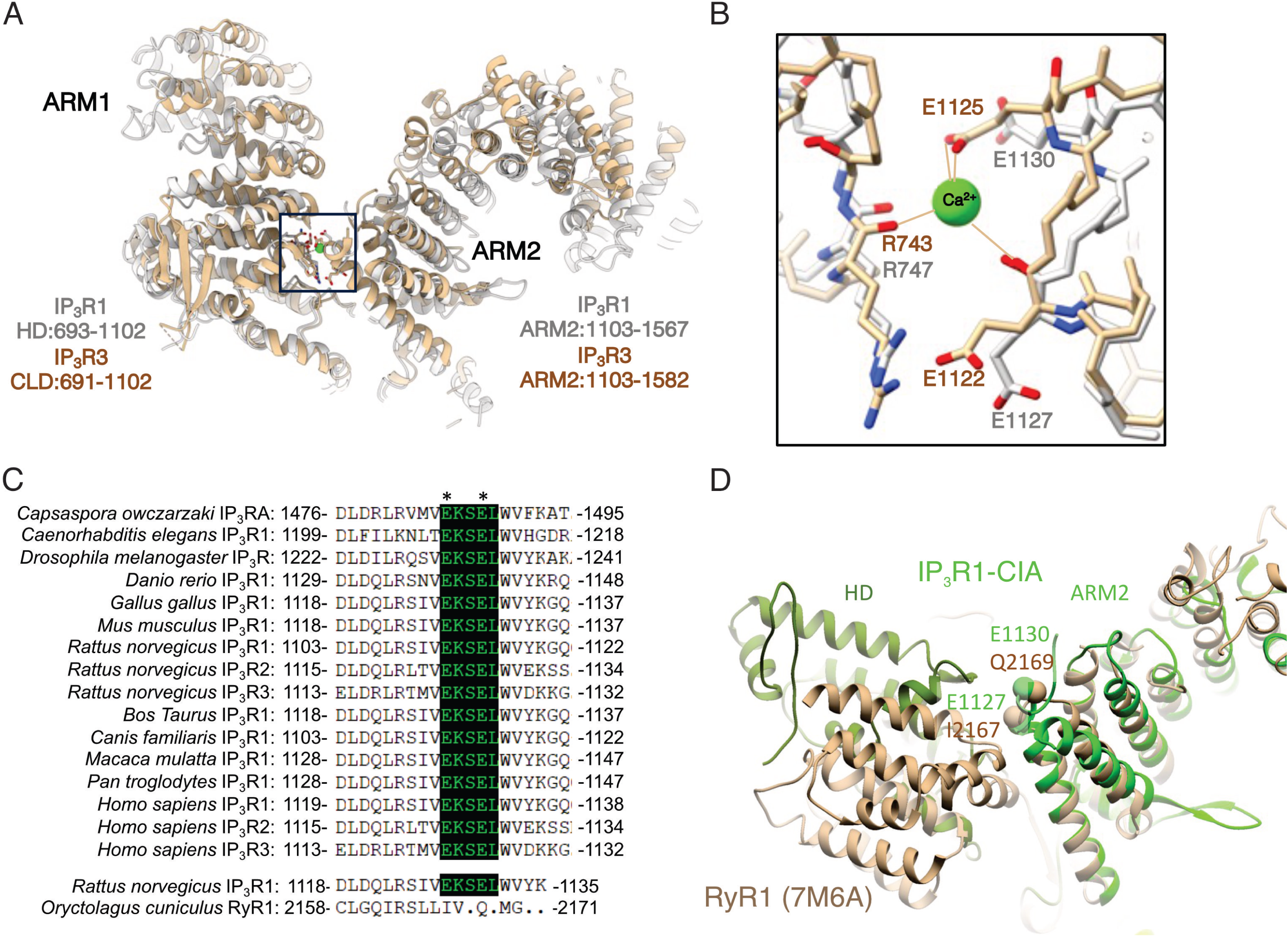
**A.** Comparison of CD Ca^2+^ binding site in IP_3_Rs. Overlay of rIP_3_R1 and hIP_3_R3 structures, highlighting the ARM2 and HD-IP_3_R1/CD-IP_3_R3 domains that form the Ca²⁺ binding site (outlined with a dashed line). **B.** Zoomed-in view of the Ca-CD site (rIP_3_R1 - 8EAR/grey; hIP_3_R3 - 6DRC/tan). The highly conserved R747-rIP_3_R1/R743-hIP_3_R3, E1127-rIP_3_R1/E1122-hIP_3_R3, E1130- rIP_3_R1/E1125-hIP_3_R3 responsible for coordination of Ca^2+^ are labeled. Notably, Ca²⁺ has only been observed in the 6DRC structure. **C.** Structural alignment of the Ca-CD site between IP_3_R subtypes and through evolution. The Ca-CD site is not conserved in RyR. **D.** Overlay of rIP_3_R1 (green) and rabbit RyR1 (tan) structures (7M6A). The IP_3_R CD Ca^2+^ binding residues are not conserved in rabbit RyR1 structure.

### Mutating the putative Ca^2+^ coordination residues in hIP_3_R1-ARM2 domain augments agonist-induced Ca^2+^ release

To determine the effects of mutating the conserved negatively charged putative Ca^2+^ coordinating E1128 and E1131 residues in hIP_3_R1-ARM2, we generated stable cell lines expressing WT IP_3_R1, mutant IP_3_R1-E1128A, and mutant IP_3_R1-E1131A constructs in HEK-3KO (HEK-IP_3_R1/2/3 triple knockout) background, previously generated in our laboratory by CRISPR-Cas9 technology (31) (Fig. 2). The E-A substitution maintains the β-carbon moiety but eliminates the negative charge on the side chain essential for Ca^2+^ coordination. To address whether these mutations would affect the overall protein architecture, a structure-based free energy difference (ΔΔG) between WT IP_3_R1 and mutant channels was calculated with ΔΔG ≈ 0, indicating that these mutations will have a negligible effect on protein stability of the tetrameric IP_3_R channel (33).

**Fig. 2.**
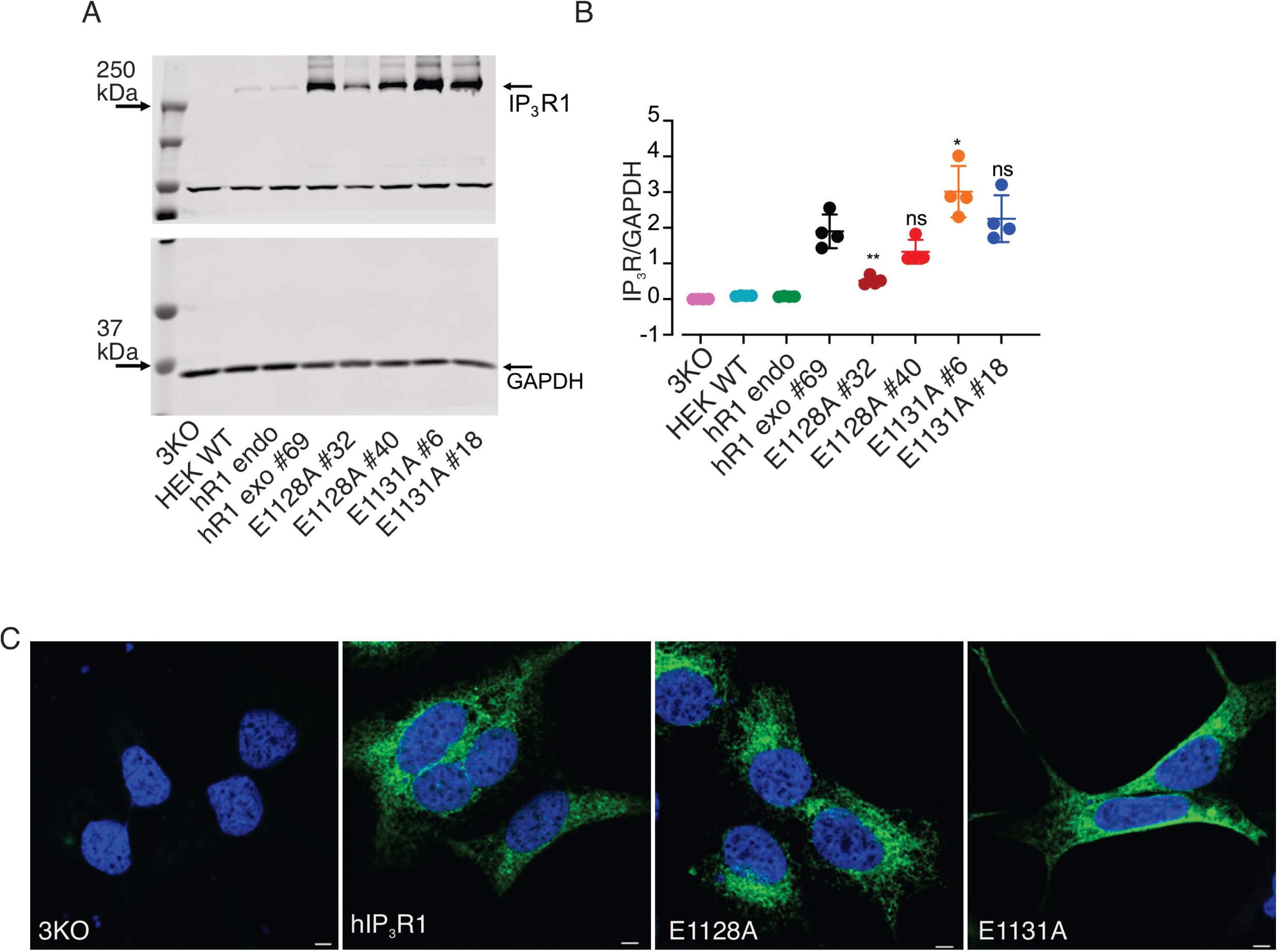
Generation of stable cells in HEK-293 IP_3_R null-background **A.** A representative western blot depicting IP_3_R1 and GAPDH protein levels in HEK-3KO, HEK WT (HEK293), hR1 endo, stable hR1 exo 69, E1128A (clones #32 and #40), and E1131A (clones #6 and #18) cell lines. **B.** Scatter plot showing quantification of IP_3_R1 protein levels normalized to GAPDH. Data are presented as mean±s.e.m from n=4 independent experiments. Statistical significance was determined by one-way ANOVA with Tukey’s test. *p<0.05, **p<0.01, ns; non-significant when compared to hR1 exo 69 cell line. **C.** Localization of IP_3_R1 in hR1 exo 69, E1128A #40, and E1131A #18 mutant stable cells. Scale bar, 10 µm.

Both WT (hR1 exo 69) and mutant stable (E1128A and E1131A) cells expressed higher IP_3_R1 protein levels when compared to HEK-293 and HEK-IP_3_R2/3 double knockout (hR1 endo) cells. Moreover, the IP_3_R1 protein levels in E1128A #40 and E1131A #18 clones were comparable to WT hR1 exo 69 clone (Figs. 2A and 2B). Therefore, all the subsequent experiments were performed using these clones. Both WT and mutant IP_3_R1 appropriately localized to the endoplasmic reticulum and displayed a distinctive reticular pattern (Fig. 2C).

Next, to determine the functional consequences of these mutations on IP_3_-induced Ca^2+^ release, we performed single-cell Ca^2+^ imaging assays in cells loaded with Fura2/AM and stimulated with carbachol (CCh), an agonist of G_α_q-coupled M3 muscarinic receptor (34). CCh-induced Ca^2+^ release was significantly potentiated in the E1128A expressing cells when compared to hR1 cells at all [CCh] tested (3 µM, 10 µM, and 30 µM) (Figs. 3A-C). The percentage of responding E1128A expressing cells was significantly higher at lower [CCh] (3 µM, and 10 µM), however, at a higher level (30 µM) of CCh, virtually all the WT and mutant cells responded (Fig. 3D). The number of hR1 and E1128A expressing cells displaying Ca^2+^ oscillations increased with rising [CCh] and surprisingly, the fraction of cells displaying Ca^2+^ oscillations were higher in the E1128A expressing cells compared to WT cells (Fig. 3E). Similarly, CCh-induced Ca^2+^ release was significantly augmented in the E1131A expressing cells when compared to hR1 cells at all [CCh] tested (0.3 µM, 1 µM, and 3 µM) (Figs. 4A-C) and the percentage of responding E1131A expressing cells was significantly higher at all [CCh] (Fig. 4D). The number of hR1 and E1131A expressing cells exhibiting Ca^2+^ oscillations increased with increasing [CCh] and similar to the E1128A expressing cells, the fraction of cells displaying Ca^2+^ oscillations were again higher in the E1131A expressing cells compared to WT cells (Fig. 4E). In addition, in population-based Ca^2+^ imaging assays performed using a FlexStation3 plate reader equipped with microfluidics, CCh-induced maximum amplitude changes were significantly enhanced in both E1128A and E1131A compared to hR1 expressing cells (Fig. 5). Overall, these results suggest that abolishing the negative charge on the E1128 and E1131 residues renders the channels more sensitive to IP_3_-generating agonists, consistent with reduced IP_3_R channel inhibition at higher [Ca^2+^]_i_ in the presence of IP_3._ These data are also in accord with the prediction that the side chain of E1125 residue in hIP_3_R3 (corresponding to E1131 residue in hIP_3_R1) and the main chain carbonyl oxygen of E1122 residue in hIP_3_R3 (corresponding to E1128 residue in hIP_3_R1) facilitate Ca^2+^ coordination to the CD site (Fig. 1C) (25, 30).

**Fig. 3.**
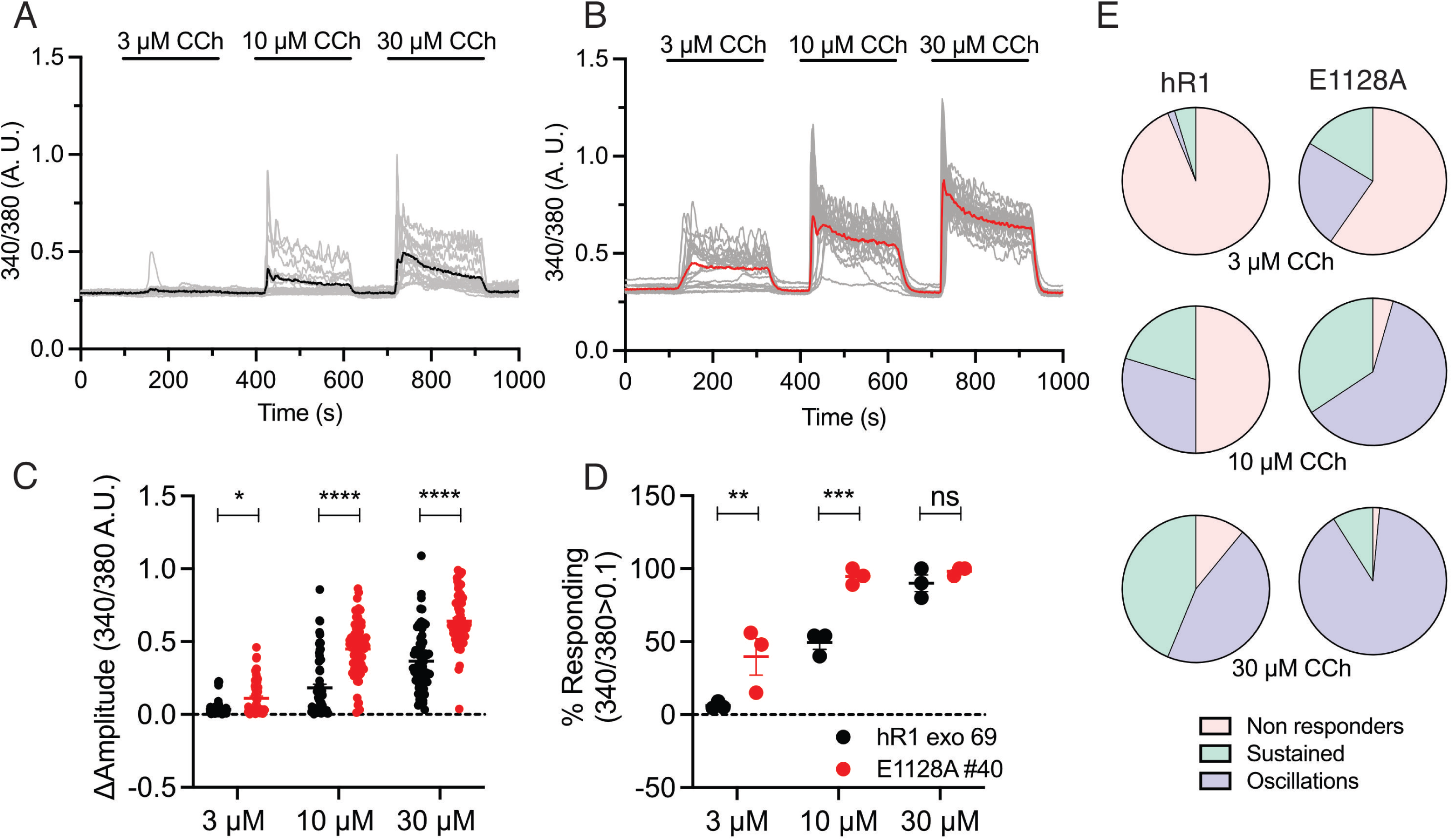
E1128A mutant stable cells exhibit enhanced sensitivity to IP_3_ generating agonist compared to WT cells. Representative Ca^2+^ traces showing increase in cytosolic Ca^2+^ levels from **A.** hR1 exo, and **B.** E1128A cells in response to stimulation with 3, 10, and 30 μM CCh. The thicker black and red traces denote the average increase in cytosolic Ca^2+^ from the population. Scatter plots comparing **C.** changes in amplitude (peak ratio – basal ratio: average of initial 20 ratio points) and **D.** percentage of responding cells (amplitude changes >0.1), from hR1 exo and E1128A cells in response to stimulation with 3, 10, and 30 μM CCh. Data are presented as mean±s.e.m from n=3 independent experiments. Statistical significance was determined by two-way ANOVA with Sidak test. *p<0.05, **p<0.01, ***p<0.001, and ****p<0.0001. ns; non-significant. **E.** Pie-charts comparing cell response heterogeneity [non-responders, sustained/global Ca^2+^ signals, and oscillating Ca^2+^ signals (amplitude changes >0.05)] between hR1 exo 69 (n=64 cells) and E1128A #40 (n=67 cells) cell lines in response to stimulation with 3, 10, and 30 μM CCh.

**Fig. 4.**
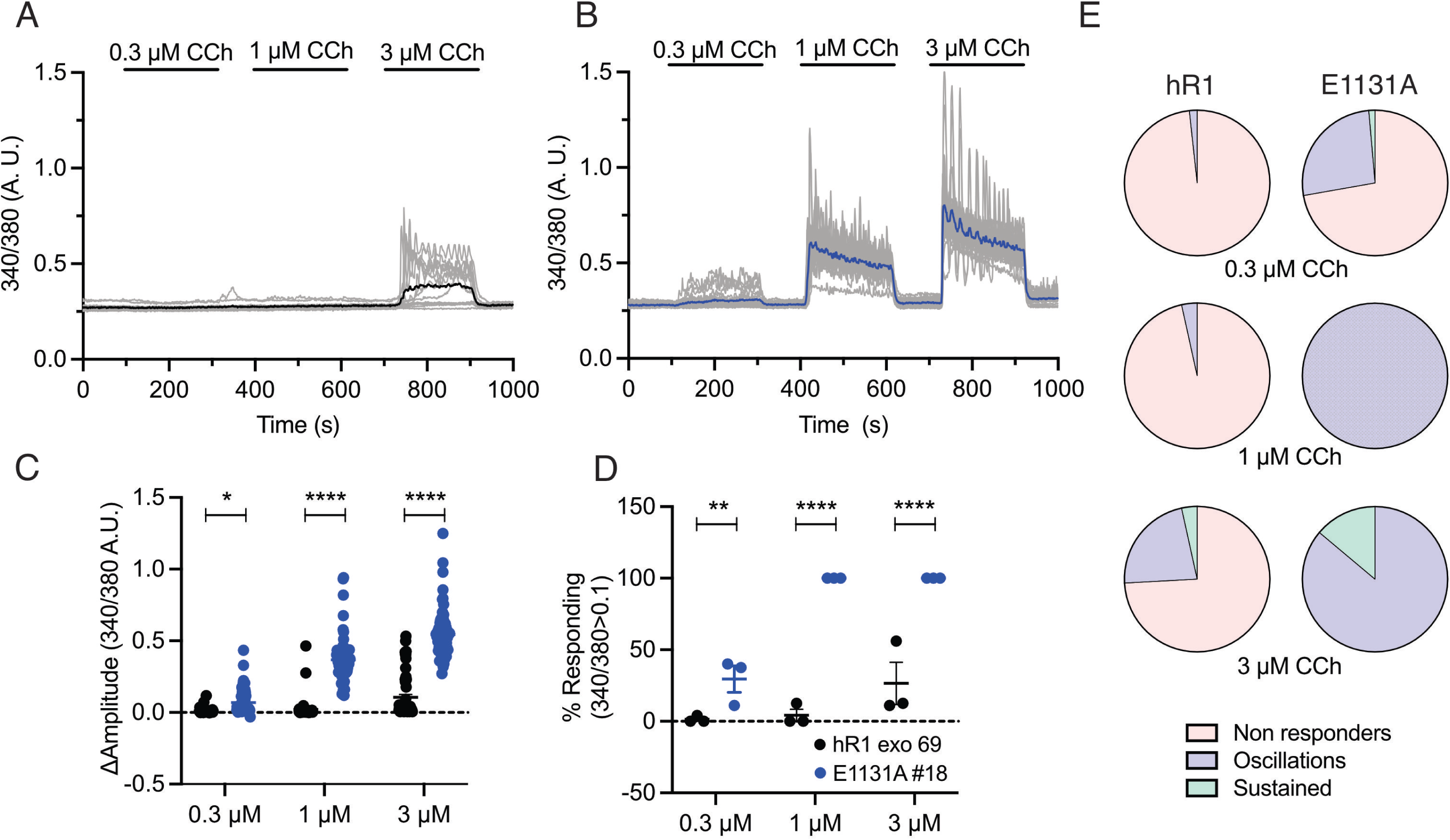
E1131A mutant stable cells exhibit enhanced sensitivity to IP_3_ generating agonist compared to WT cells. Representative Ca^2+^ traces showing increase in cytosolic Ca^2+^ levels from **A.** hR1 exo and **B.** E1131A cells in response to stimulation with 0.3, 1, and 3 μM CCh. The thicker black and blue traces denote the average increase in cytosolic Ca^2+^ from the population. Scatter plots comparing **C.** changes in amplitude (peak ratio – basal ratio: average of initial 20 ratio points) and **D.** percentage of responding cells (amplitude changes >0.1), from hR1 exo and E1131A cells in response to stimulation with 0.3, 1, and 3 μM CCh. Data are presented as mean±s.e.m from n=3 independent experiments. Statistical significance was determined by two-way ANOVA with Sidak test. *p<0.05, **p<0.01, and ****p<0.0001. **E.** Pie-charts comparing cell response heterogeneity [non-responders, sustained/global Ca^2+^ signals, and oscillating Ca^2+^ signals (amplitude changes >0.05)] between hR1 exo (n=58 cells) and E1131A (n=72 cells) cell lines in response to stimulation with 0.3, 1, and 3 μM CCh.

**Fig. 5.**
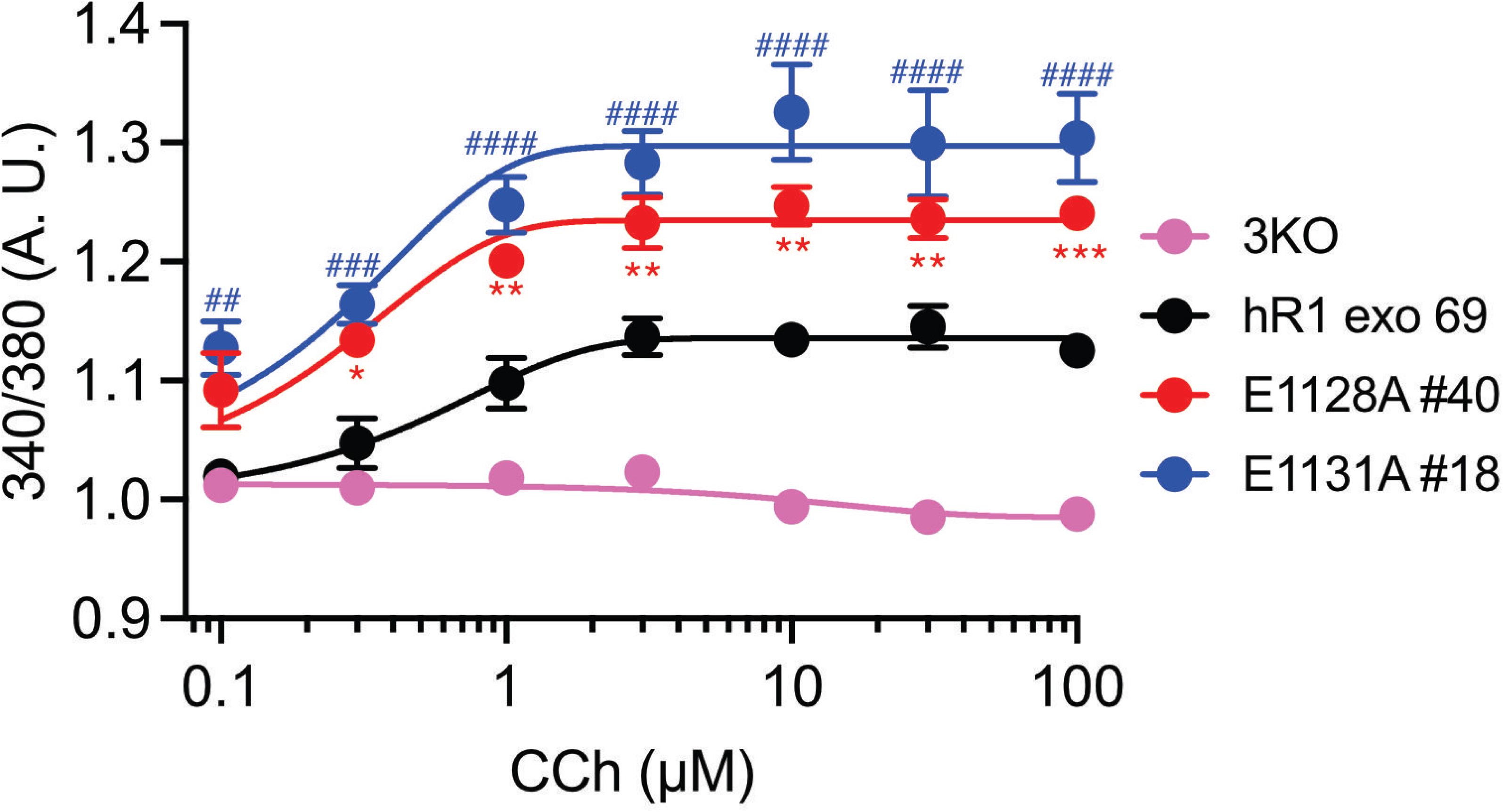
CD site Ca^2+^ binding mutants displayed augmented Ca^2+^ release in population-based Ca^2+^ imaging assays. Dose-response curves showing Ca^2+^ responses from Fura-2/AM loaded HEK-3KO, hR1 exo, E1128A, and E1131A cell lines when treated with increasing concentrations (0.1 μM, 0.3 μM, 1 μM, 3 μM, 10 μM, 30 μM, and 100 μM) of CCh (n=3 independent experiments). Data are presented as mean±s.e.m from n=3 independent experiments. Statistical significance was determined by two-way ANOVA with Tukey’s test. *p<0.05, **p<0.01, and ***p<0.001 between hR1 exo vs E1128A cells. ##p<0.01, ###p<0.001, and ####p<0.0001 between hR1 exo vs E1131A cells.

### The CD site Ca^2+^ binding mutants exhibit spontaneous Ca^2+^ puffs and are more sensitive to IP_3_ stimulation

To directly measure the elementary Ca^2+^ signals from IP_3_Rs (called Ca^2+^ puffs), we used TIRF microscopy (35, 36). Surprisingly, under basal, resting conditions without any additional overt stimulation, spontaneous Ca^2+^ puffs were readily evident in both E1128A and E1131A, but absent in WT expressing cells, as shown in maximal-intensity projection images binned over time, representative traces, and pooled data (Figs. 6A-C). These results suggest that the removal of negative charges in the Ca-CD site renders the channel more sensitive to basal ambient levels of IP_3_. Another remote possibility is that the channel becomes more susceptible to spontaneous opening in the absence of IP_3_. The latter option was deemed unlikely because no channel activity was observed in the absence of pipette IP_3_ in “on-nucleus“ patch clamp experiments in DT40-3KO cells stably expressing E1128A and E1131A, or in cells expressing a double mutation of both residues (Fig. 6D).

**Fig. 6.**
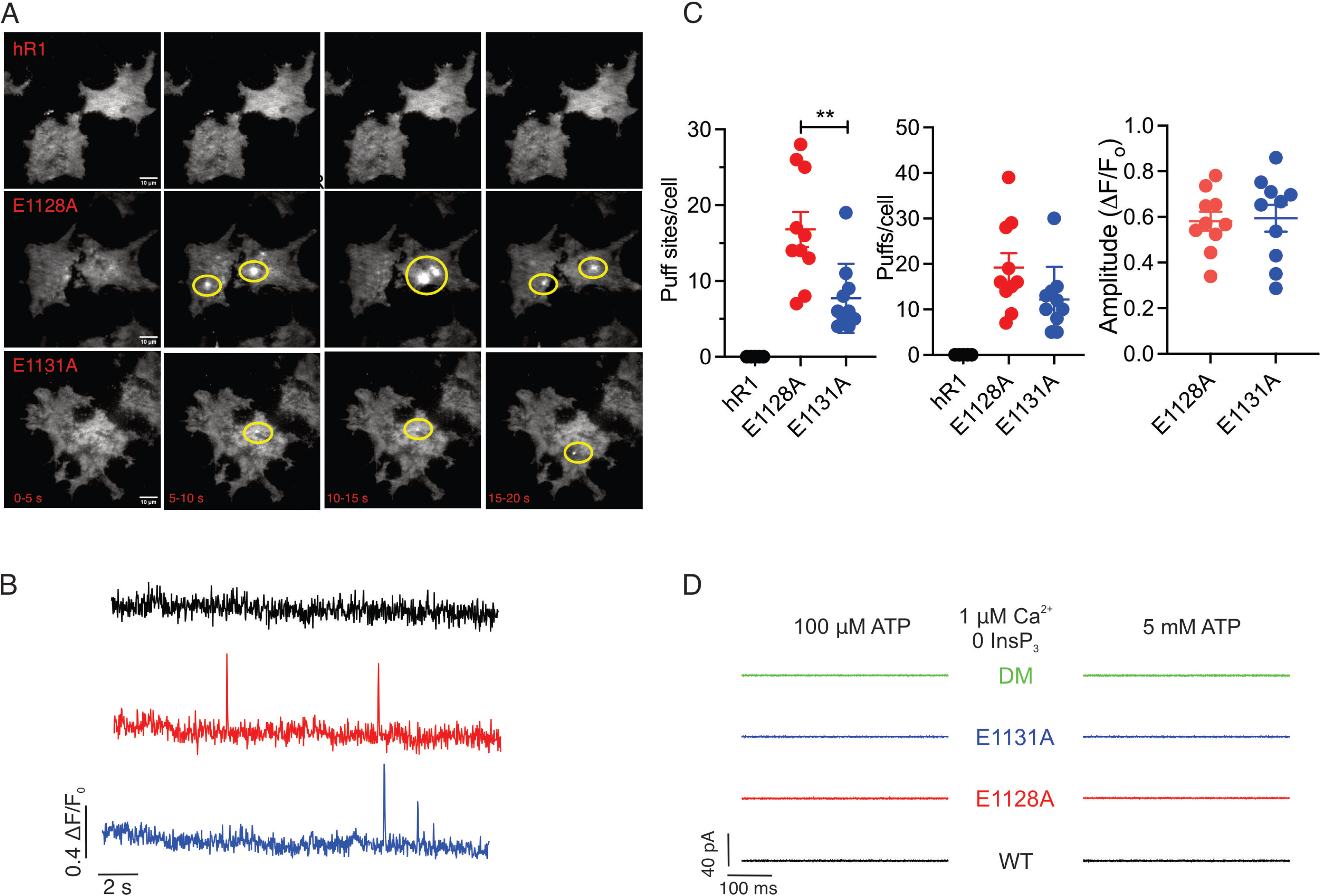
CD site Ca^2+^ binding mutants exhibit spontaneous Ca^2+^ puffs under basal conditions. **A.** Maximal intensity projections of Cal-520 fluorescence at 5 s intervals from hR1 exo, E1128A, and E1131A cell lines under basal resting conditions. Elementary Ca^2+^ signals in E1128A, and E1131A cells are highlighted. **B.** Representative traces showing fluorescence changes from the center of a single puff site (1 × 1 μm) from hR1 exo (black), E1128A (red), and E1131A (blue) cell lines. **C.** Scatter plots summarizing number of puff sites/cell, number of puffs/cell, and average amplitudes of the Ca^2+^ puffs are shown for hR1 exo, E1128A, and E1131A cells. Data are presented as mean±s.e.m from n=10 independent experiments. Statistical significance was determined by one-way ANOVA with Tukey’s test. **p<0.01. **D.** Representative sweeps obtained using “on-nucleus” patch clamp experiments from DT-40 3KO cells stably expressing either WT (R1) or mutant (E1128A, E1131A or CD DM) constructs without IP_3_ in the presence of either 100 µM or 5 mM ATP and 1 µM [Ca^2+^].

We next increased IP_3_ by photolysis of a threshold concentration of caged inositol triphosphate (ci-IP_3_) (cell loaded with 0.05 µM of ci-IP_3_/PM) to capture IP_3_-induced Ca^2+^ puffs. As shown in the representative traces and maximal-intensity projections binned over time, the number of Ca^2+^ signals from the E1128A and E1131A expressing cells were augmented compared to the hR1 expressing cells (Figs. 7A and 7E). The number of puffs per cell, and puff sites per cell were significantly higher in the E1128A and E1131A expressing cells compared to hR1 expressing cells (Figs. 7B and 7C). The amplitudes and rise times did not differ between the WT and mutant cells; however, the fall/decay times were significantly longer for the mutant cells compared to the WT cells (Figs. 7D and 7F), consistent with reduced Ca^2+^ dependent inhibition of IP_3_R activity. These results indicate Ca^2+^ binding to the CD site may be critical to facilitate IP_3_R channel closure.

**Fig. 7.**
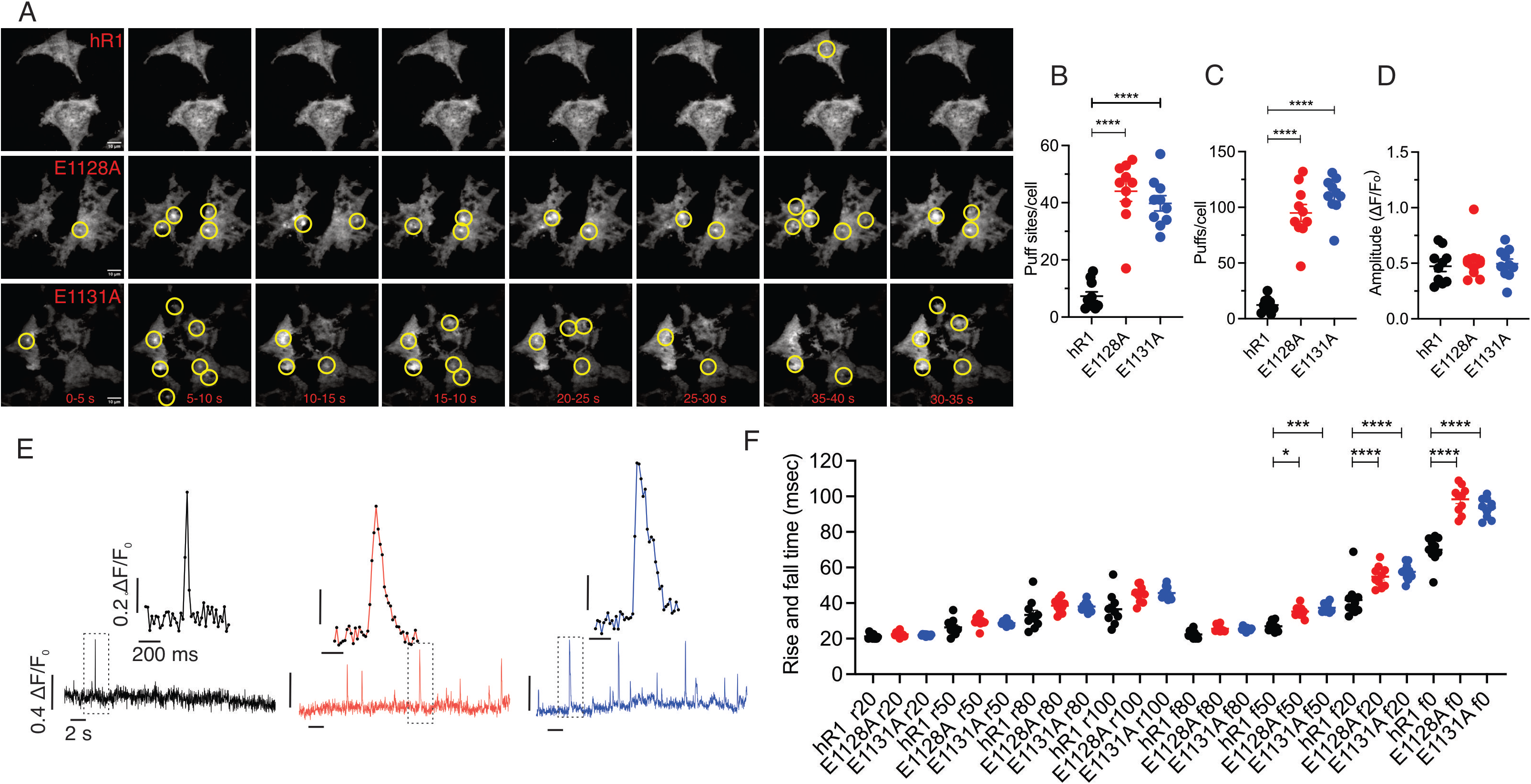
CD site Ca^2+^ binding mutants exhibit augmented Ca^2+^ puffs upon uncaging ci-IP_3_. **A.** Maximal intensity projections of Cal-520 fluorescence at 5 s intervals from hR1 exo, E1128A, and E1131A cell lines upon uncaging 0.05 µM ci-IP_3_. Elementary Ca^2+^ signals are highlighted. **E.** Representative traces showing fluorescence changes upon uncaging ci-IP_3_ from the center of a single puff site (1 × 1 μm) from hR1 exo (black), E1128A (red), and E1131A (blue) cell lines. Individual Ca^2+^ puffs from the boxed area highlighting longer decay/fall time for the CD site Ca^2+^ binding mutants when compared to hR1 exo cells. Scatter plots summarizing **B.** number of puff sites/cell, **C.** number of puffs/cell, **D.** average amplitudes of the Ca^2+^ puffs, and **F.** mean-rise (r) and -fall (f) times for the fluorescence to rise/decay to various levels (0%–100%) are shown. Data are presented as mean±s.e.m from n=10 independent experiments. Statistical significance was determined by one-way ANOVA with Tukey’s test. **p<0.01, **p<0.01, ***p<0.001, and ****p<0.0001.

### CD site Ca^2+^ binding double mutant is more sensitive to IP_3_

We generated stable WT IP_3_R1, mutant IP_3_R1-E1128A, mutant IP_3_R1-E1131A, and IP_3_R1-E1128A/E1131A double mutant (CD-DM) in DT40-3KO (IP_3_R1/2/3 triple knockout) background (Figs. 8A and 8B) (32). We performed single-cell Ca^2+^ imaging assays in cells loaded with Fura2/AM and varying amounts of ci-IP_3_. As shown in Fig. 8C, DT40 CD-DM cells displayed augmented Ca^2+^ signals in response to repeated UV-mediated flash photolysis of low concentration of ci-IP_3_ (cell loaded with 1 µM ci-IP_3_/PM) while the hR1 expressing cells were essentially non-responsive. The hR1 expressing cells responded to uncaging higher levels of ci-IP_3_ (5 µM) although the Ca^2+^ signals from these cells were attenuated when compared to DT40 CD-DM cells (Fig. 8D). These results confirm that the double mutant is functional, and like the E1128A and E1131A mutant expressing HEK cells, exhibits augmented IP_3_-induced Ca^2+^ release consistent with reduced inhibitory input to channel activity.

**Fig. 8.**
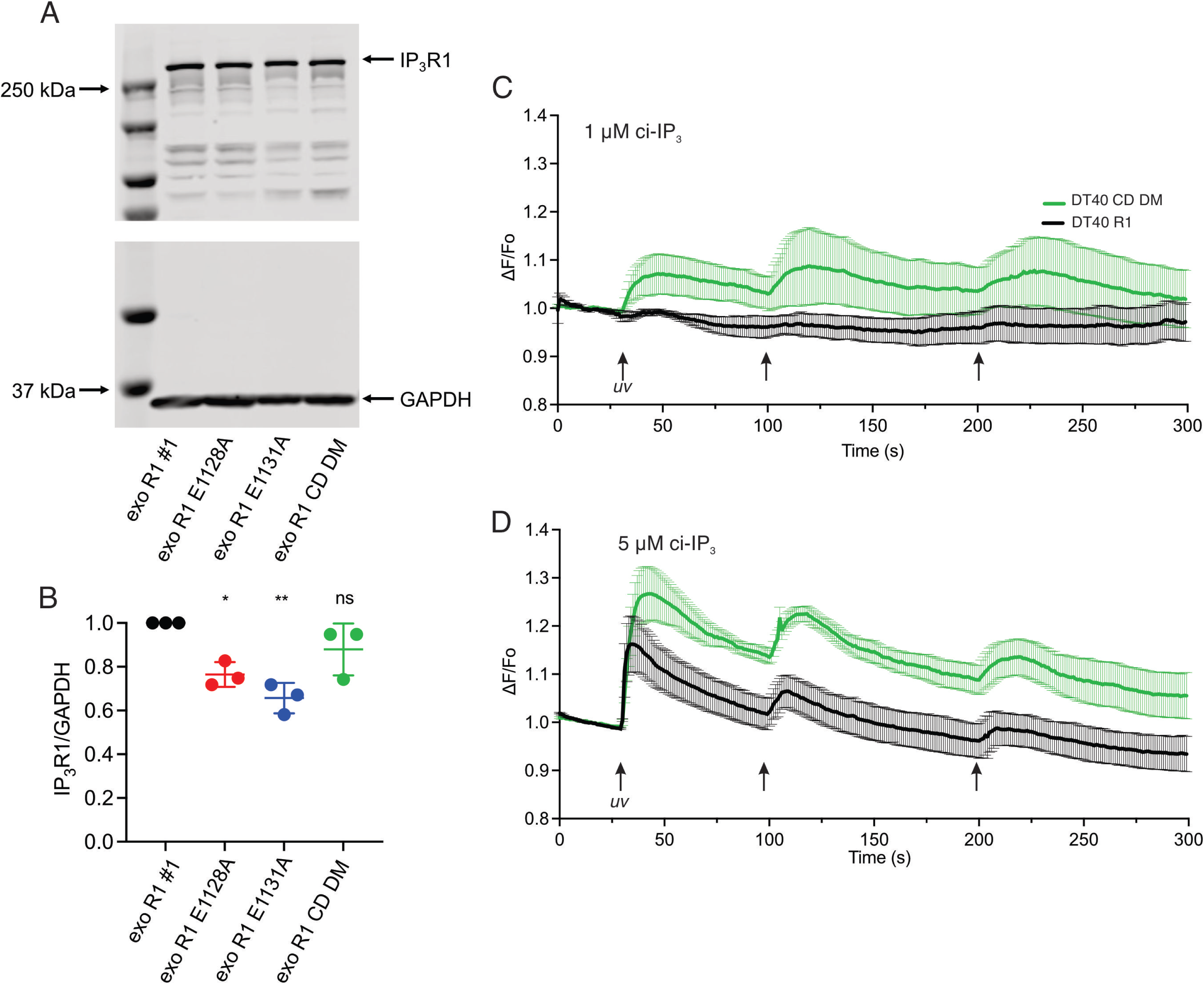
Generation of stable cells in DT-40 3KO background. **A**. A representative western blot depicting IP_3_R1 and GAPDH protein levels in DT40 exo R1 (clone #1), DT40 exo R1 E1128A, DT40 exo R1 E1131A, and DT40 exo R1 CD DM (E1128/1131A) cell lines. **B.** Scatter plot showing quantification of IP_3_R1 protein levels normalized to GAPDH. Data are presented as mean±s.e.m from n=3 independent experiments. Statistical significance was determined by one-way ANOVA with Tukey’s test. *p<0.05, **p<0.01, ns; non-significant when compared to DT40 exo R1 cell line. A comparison of intracellular global Ca^2+^ signals (ΔF/F_o_) between WT (DT40 R1) and CD DM cells following repeated photolysis of either **C.** 1 µM or **D.** 5 µM ci-IP_3_ at indicated time points (arrows). Data are presented as mean±s.e.m from n=3 independent experiments.

### The CD site is not responsible for IP_3_R channel closure at saturating free ATP

We next performed “on-nucleus” patch-clamp experiments in DT-40 cells stably expressing the WT (R1), single (E1128A and E1131A), and double mutants (CD-DMs) in the presence of sub maximal ([IP_3_]; 1 µM), saturating free ATP ([Na-ATP]; 5 mM), and varying [Ca^2+^]. The open probability of the R1 channels increased with increasing [Ca^2+^] and peaked at ∼0.2 µM [Ca^2+^] (∼0.27). A further increase in [Ca^2+^], diminished the open probability of the R1 channels and channels were essentially closed at 100 μM [Ca^2+^] displaying a characteristic bell-shaped Ca^2+^-mediated regulation of IP_3_R1 channel activity (Fig. 9A). Surprisingly, both single and the DM channels exhibited a similar dependency on Ca^2+^ for activation and inhibition (Figs. 9A and 9B). These results indicate that Ca^2+^ binding to the CD site does not facilitate IP_3_R channel closure at saturating [ATP].

**Fig. 9.**
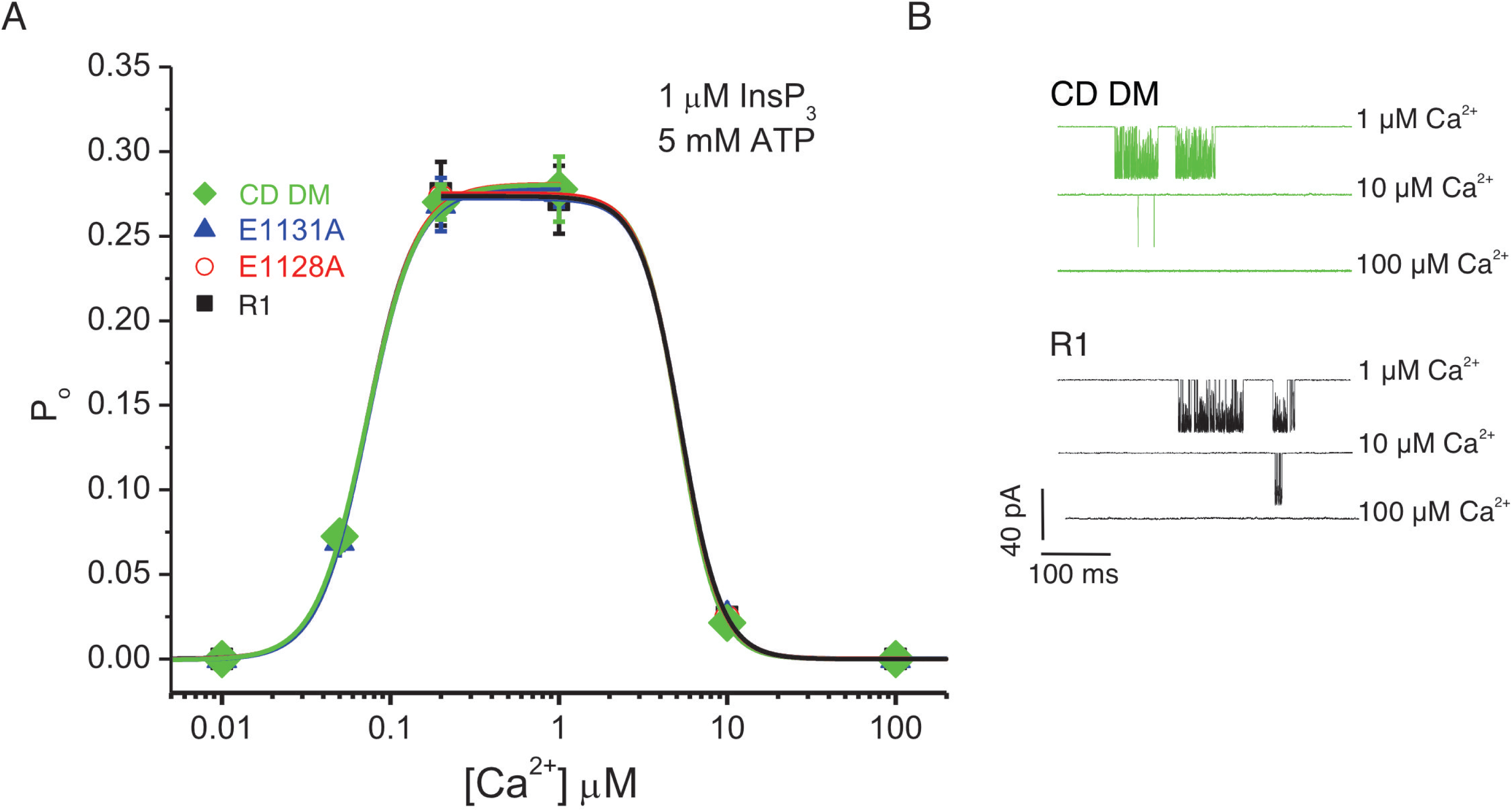
CD site is not essential for Ca^2+^-dependent inactivation of IP_3_R1 at high [ATP]. **A.** Hill equation fit for pooled data obtained using “on-nucleus” patch clamp experiments from DT-40 3KO cells stably expressing either WT (R1) or mutant (E1128A, E1131A or CD DM) constructs at indicated [Ca^2+^] stimulated with 1 µM IP_3_ in the presence of 5 mM ATP. **B.** Representative sweeps for the CD DM and hR1 at indicated [Ca^2+^] in the presence of 1 µM IP_3_ and 5 mM ATP are shown.

### The CD site is essential for IP_3_R channel inactivation at sub-saturating [ATP]

Since previous results from our group and others established that ATP potentiates IP_3_R1 channel activity by tuning the sensitivity of IP_3_R to Ca^2+^ (15, 16, 37), we next performed experiments at 100 μM [ATP], close to the EC_50_ value (150 μM) determined in permeabilized DT40-3KO cells expressing rIP_3_R1 (15, 38). Notably, the open probability of the R1 and mutant channels activated by 1 μM IP_3_ displayed a similar Ca^2+^ dependency for channel activation (P_o_∼0.25 at 1 and 10 µM [Ca^2+^]) (Fig. 10A). Nevertheless, the mutants exhibited a significantly higher P_o_ at inhibitory [Ca^2+^] (>10 µM [Ca^2+^]) when compared to the WT channels (Figs. 10A and 10B). The DM and E1131A mutant channels displayed a higher P_o_, at elevated [Ca^2+^] indicating less inhibition by high [Ca^2+^] compared to E1128A mutant channels. These data are consistent with the increased CCh-induced Ca^2+^ release in E1131A vs E1128A expressing cells (Figs. 3, 4, and 5). High Ca^2+^-induced inhibition of channel activity was similarly reduced in DM mutant channels when the [IP_3_] was increased to 10 μM (Fig. 10C and 10D). In total these data indicate that high Ca^2+^-induced inhibition of IP_3_R channel activity is mediated via the CD Ca^2+^ binding site only at sub-saturating [ATP].

**Fig. 10.**
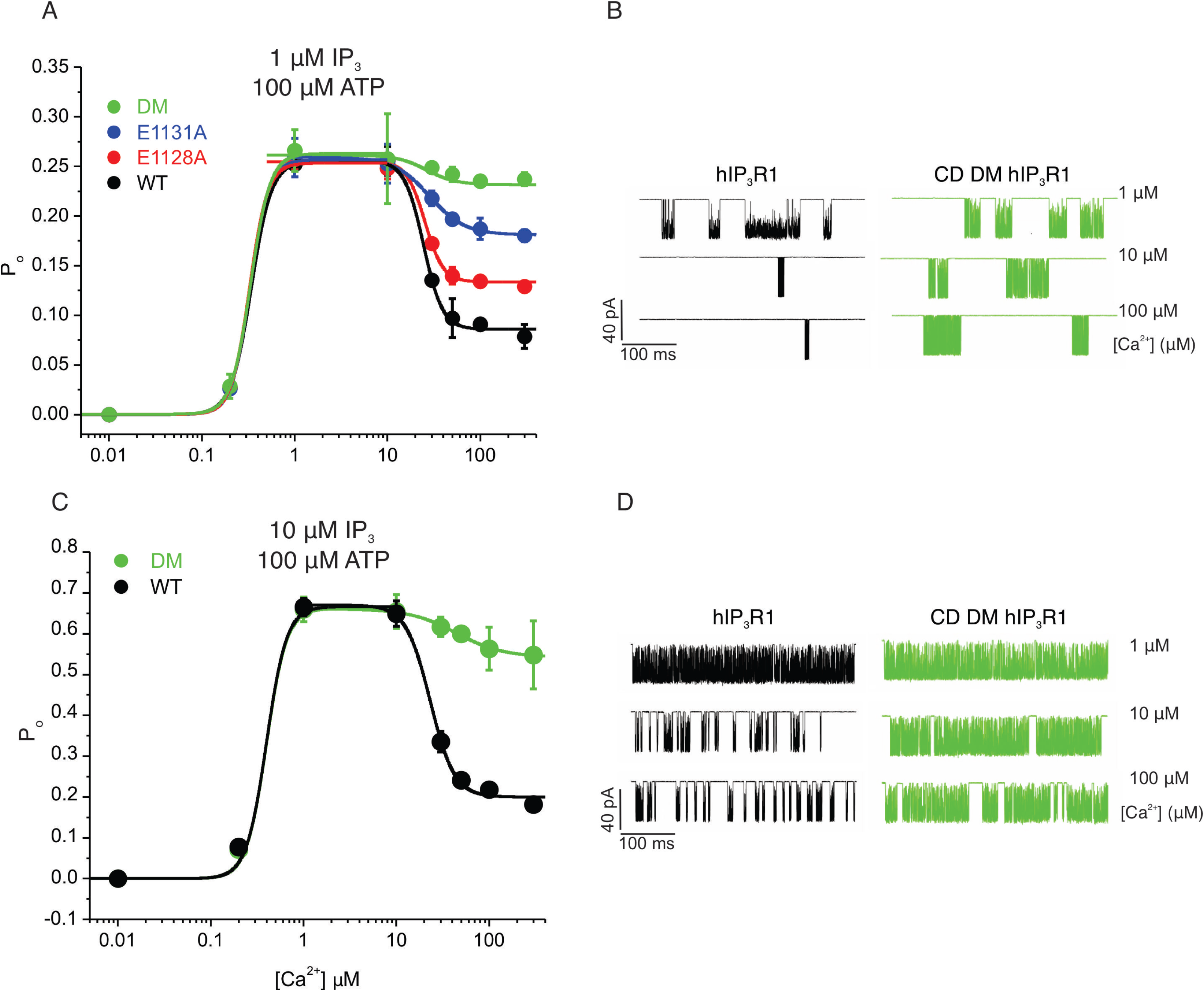
CD site is critical for Ca^2+^-dependent inactivation of IP_3_R1 at low [IP_3_] and [ATP]. **A.** Hill equation fit for pooled data obtained using “on-nucleus” patch clamp experiments from DT-40 3KO cells stably expressing either WT (R1) or mutant (E1128A, E1131A or CD DM) constructs at indicated [Ca^2+^] stimulated with 1 µM IP_3_ in the presence of 100 µM ATP. **B.** Representative sweeps showing higher activity from CD DM compared to R1 at inhibitory [Ca^2+^]. **C.** Hill equation fit for pooled data obtained using “on-nucleus” patch clamp experiments from DT-40 3KO cells stably expressing either WT (R1) or CD DM mutant constructs at indicated [Ca^2+^] stimulated with 10 µM IP_3_ in the presence of 100 µM ATP. **D.** Representative sweeps showing higher activity from CD DM compared to R1 at inhibitory [Ca^2+^].

## Discussion

The activity of IP_3_Rs is determined by their endogenous primary ligands IP_3_ and Ca^2+^ (2). It is well established that [Ca^2+^]_i_ regulates IP_3_R channel activity in a biphasic manner by binding to intrinsic non-canonical Ca^2+^ binding sites within the receptor (10, 11). Free ATP (free ATP^4-^) allosterically modulates IP_3_R activity by enhancing the sensitivity of IP_3_Rs to Ca^2+^ (9, 39–41), resulting in a shift in Ca^2+^ activation and inhibition of IP_3_R. Based on the available cryo-EM structures of IP_3_Rs, we previously identified evolutionarily well-conserved Ca^2+^ binding residues critical to facilitate IP_3_R channel activation (27). Mutating the Ca^2+^ activation site (Ca-III_s_/JD site) did not perturb Ca^2+^-dependent channel inactivation signifying that one or more of the other putative Ca^2+^ binding sites identified in the cryo-EM structures could serve as the Ca^2+^ inhibitory site/s (24, 25, 27). Two Ca^2+^ binding sites previously identified in the cryo-EM structures of rIP_3_R1, Ca-I_LBD_ and Ca-II_LBD_ are located in the ligand binding domain while the other two Ca^2+^ binding sites, Ca-IV_L_ and Ca-V_L_, are within the luminal side of the ion conduction pathway (24). A mutational analysis approach may not be appropriate for these sites as altering these residues would be predicted to grossly alter IP_3_ binding affinity and ion permeability, respectively. Moreover, single-channel recordings from excised IP_3_Rs in luminal-side-out configuration suggest that the luminal Ca^2+^ binding sites do not functionally regulate channel activity (42). These observations indicate that Ca-IV_L_ and Ca-V_L_ represent high-affinity Ca^2+^ binding sites in the channel conduction pathway (24). The putative CD Ca^2+^ binding site was identified in the cryo-EM structure of hIP_3_R3 bound to Ca^2+^ only in the presence of IP_3_, ATP, and high [Ca^2+^]_i_ (30). Consistent with the requirement for concurrent IP_3_, ATP, and high [Ca^2+^] binding, Ca^2+^ binding to the CD site was not observed in the rIP_3_R1 structure resolved in the presence of inhibitory [Ca^2+^] without IP_3_ and ATP (24). These observations indicate that IP_3_ and Ca^2+^-induced conformational motions in the intrinsically flexible ARM2 domain facilitate CD site formation (30).

Since the putative CD site is highly-conserved in IP_3_Rs and occupied in the presence of inhibitory [Ca^2+^], we investigated the functional consequences of mutating these residues. We engineered charge mutations in the hIP_3_R1 CD Ca^2+^ binding site and generated stable cells in the HEK-3KO and DT40-3KO background, with expression levels comparable to the WT hIP_3_R1. We confirmed the mutant channels like the WT channels localized properly to the ER. Moreover, the mutant cells responded to an IP_3_-generating agonist indicating that the mutant channels can form functional homotetramers. In single cell, population-based, and TIRFM Ca^2+^ imaging assays, the mutant stable cells exhibited higher sensitivities to stimulation when compared to the WT cells. We were unsuccessful in generating a stable double mutant line in HEK-3KO cells, but a double mutant in the DT40-3KO background was also more responsive to activation by IP_3_ when compared to the WT cells.

TIRFM imaging revealed that cells expressing CD mutant IP_3_R exhibited Ca^2+^ puffs in the absence of exogenous stimulation of the cells (36). Because IP_3_ is required to activate the CD binding site mutants, this also likely reflects the increased sensitivity of CD IP_3_R mutant to IP_3_, in which these cells respond to basal endogenous levels of IP_3_. Interestingly, the heterozygous *ITPR3*-T1424M mutation in the ARM2 domain, associated with neuropathy (43, 44), when stably expressed in HEK-3KO cells also exhibited spontaneous Ca^2+^ puffs (45). Moreover, the T1424M mutant channels exhibited increased open probability at lower [IP_3_] and remained active even at inhibitory [Ca^2+^] (44). These observations underscore a critical role of ARM2 residues and the associated structural transitions between the extended and retracted ARM2 conformations in proper inactivation of IP_3_Rs at inhibitory [Ca^2+^] (24, 30).

In a previous study, mutation of the analogous residues in hIP_3_R3 resulted in a decrease in the frequency of Ca^2+^ oscillations stimulated by CCh compared to cells expressing WT hIP_3_R3, consistent with the CD site contributing to the mechanism underlying oscillatory activity (30). We noted, however, that cells expressing CD mutants had a greater propensity to exhibit Ca^2+^ oscillations. The reasons for this discrepancy are not clear, however, in the previous study, GFP-tagged IP_3_R3 constructs were transiently expressed by viral transduction in HEK-3KO cells, and the expression level of the constructs was not reported, leaving the possibility that variable expression level may have contributed to these data. An increase in oscillatory activity in our study indicates that the predominant mechanism in HEK cells likely reflects oscillating IP_3_ levels, so-called type-2 oscillations, rather than through feedback processes intrinsic to the IP_3_R1.

Next, in order to directly test the effect of mutating the CD site Ca^2+^ coordinating residues on single channels, we performed “on-nucleus” patch-clamp experiments using DT-40 stable cells expressing the WT and mutant IP_3_R1 channels. The mutant channels like the WT channels had a similar biphasic dependency on Ca^2+^ for both channel activation and inhibition at saturating [ATP]. ATP is an allosteric modulator of IP_3_R channel activity, and augments the sensitivity of IP_3_Rs to Ca^2+^, therefore, we performed these assays at 100 μM [ATP] which was close to the EC_50_ for IP_3_-induced Ca^2+^ release, reported previously in permeabilized cells (150 μM) (9, 15–17). We also note that while the level of the major cellular form of ATP, Mg-ATP is in the millimolar range, IP_3_R are regulated by free ATP, and the cytosolic levels of ATP^4-^are estimated to be 10-100-fold less and these levels are physiologically/pathologically relevant. At lower [ATP], the open probability of mutant channels was significantly higher at inhibitory [Ca^2+^] in comparison to WT channels without altering the Ca^2+^ sensitivity for activation. Overall, our results indicate that, in the presence of IP_3_, at sub saturating [ATP], Ca^2+^ binding to the Ca-III_S_/JD site promotes IP_3_R channel opening while Ca^2+^ binding to the CD site is critical for channel closure. We speculate, at sub saturating cellular [ATP], Ca^2+^ binding to the CD site may promote cell survival by preventing [Ca^2+^]_i_ overload and permit energy conservation by reducing ATP consumption by plasma membrane Ca^2+^ ATPase (PMCA) and sarcoendoplasmic reticulum Ca^2+^ ATPase (SERCA) pumps. Higher [ATP] may induce additional conformational changes in IP_3_R1 structure exposing an alternate distinct low affinity Ca^2+^ binding site, perhaps the Ca-I_LBD_ site formed by β-TF1 and β-TF2 domains of adjacent subunits, facilitating channel inactivation (24).

Notably, all three IP_3_R sub-types are enriched at specialized junctions between the ER and mitochondria in various tissues/cell types (46). At these junctions, IP_3_Rs deliver high concentrations of Ca^2+^ to the low affinity mitochondrial Ca^2+^ uptake machinery (47, 48). Such an arrangement may also aid precise tuning of IP_3_R-mediated Ca^2+^ release dictated by rapid changes in [ATP] for reliable inter-organellar communication between ER and mitochondria. Overall, our results reveal ATP, in a concentration-dependent fashion, may cause subtle conformational changes in IP_3_R structure which regulates Ca^2+^-dependent reduction in IP_3_R channel activity. Future investigations mutating the other putative Ca^2+^ binding sites are necessary to delineate the molecular mechanisms of IP_3_R regulation by ATP (24). Moreover, since IP_3_R tetramers are assembled from four homo-/hetero-monomers, the stoichiometry of CD site Ca^2+^ binding for IP_3_R channel inhibition remains to be determined (2). Concatenated receptors with one or more CD Ca^2+^ binding sites mutated will be used to address this issue (49).

## Materials and Methods

### Alignment of IP_3_R protein sequences from various organisms

Alignment of IP_3_R protein sequences from *Capsaspora owczarzaki*-IP_3_RA (XP_004347577.1); *Caenorhabditis elegans*-IP_3_R1 (NP_001023170.1); *Drosophila melanogaster*-IP_3_R (NP_730941.1); *Danio rerio*-IP_3_R1 (XP_021335554.1); *Gallus gallus*-IP_3_R1 (NP_001167530.1); *Mus musculus*-IP_3_R1 (NP_034715.3); *Rattus norvigecus*-IP_3_R1 (NP_001257525.1), IP_3_R2 (NP_112308.1), and IP_3_R3 (NP_037270.2); *Bos taurus*-IP_3_R1 (NP_777266.1); *Canis familiaris*-IP_3_R1 (XP_005632286.1); *Macaca mulatta*-IP_3_R1 (NP_034715.3); *Pan troglodytes*-IP_3_R1 (XP_009443057.1); and *Homo sapiens*-IP_3_R1 (NP_001093422.2), IP_3_R2 (NP_002214.2), IP_3_R3 (NP_002215.2) were generated using GeneDoc. The cryo-EM structure of rat-IP_3_R1 (NP_P29994.2) was compared with rabbit-RyR1 (NP_001095188.1).

### Calculation of protein stability

Changes in free energy differences (ΔΔG) were carried out on the IP_3_R1 cryo-EM structures, 8EAR and 7LHE, for all mutations generated in this study using MAESTRO (33) and DDGun (50). All ΔΔGs calculated were between −0.32 and −0.04, with a predicted confidence greater than 0.77, indicating that the mutations are highly likely to not destabilize the protein folding.

### Plasmid constructs

The glutamic acid (E) residue at position 1128 and 1131 in hIP_3_R1 (NP_001093422.2) open reading frame cloned in pDNA3.1 expression plasmid were each mutated to alanine using Q5 site-directed mutagenesis kit (NEB) and appropriate primers obtained from Integrated DNA technologies (Table 1). In order to generate the double mutant (E1128A/E1131A), E1128A mutation harboring plasmid was used as a template to mutate the E1131 residue to alanine using specific primers (Table 1). The successful incorporation of desired mutations were confirmed by Sanger sequencing.

**Table 1:**
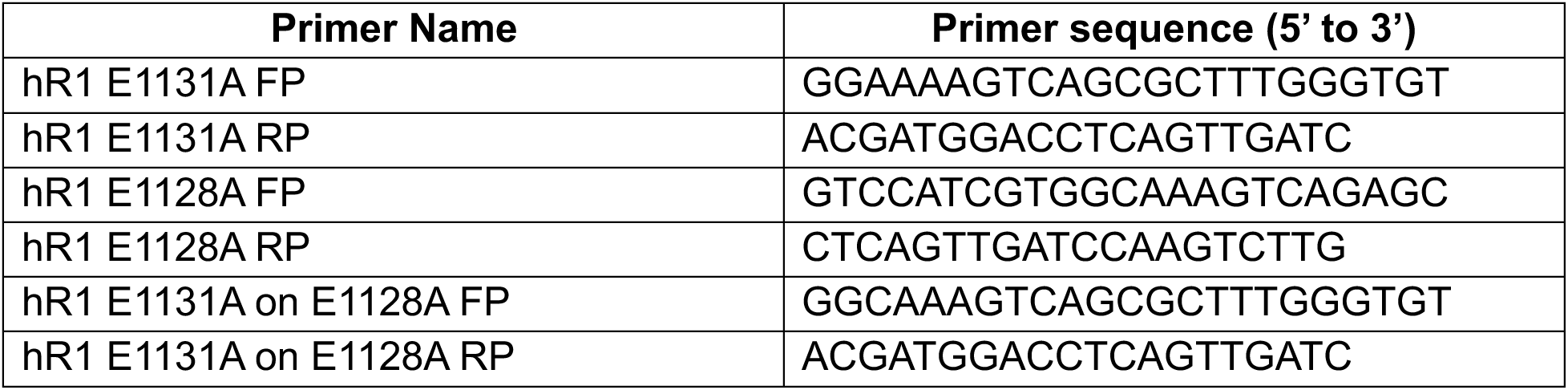
Primers used to generate substitutions at the desired site.

### Cell culture, transfection, and generation of stable cell lines

DT40 cells, chicken B lymphocyte, engineered to lack all the three native IP_3_R sub-types (DT40-3KO) were obtained (32). DT40-3KO cells were cultured in RPMI 1640 media supplemented with 1% chicken serum, 10% fetal bovine serum, 100 U/mL penicillin, and 100 μg/mL streptomycin in an incubator set to 39°C with 5% CO_2_. DT40-3KO cells were transfected as previously described (51). Briefly, 5 million cells were washed with phosphate-buffered saline (PBS) and electroporated with 4 to 6 μg of WT IP_3_R1, E1128A, E1131A or E1128A/E1131A plasmid construct using an Amaxa cell nucleofector (Lonza Laboratories) and nucleofection reagent (362.88 mM ATP-disodium salt, 590.26 mM MgCl_2_ 6.H2O, 146.97 mM KH_2_PO_4_, 23.81 mM NaHCO_3_, and 3.7 mM glucose at pH 7.4). The cells were allowed to recover for 24 h and were subsequently transferred into 96-well plates containing media supplemented with 2 mg/mL G418 (#11811-031, Gibco). Next, 10 to 14 days after transfection, clones expressing the desired construct were expanded and subsequently screened by western blotting. Cell lines stably expressing the construct were used in further experiments.

HEK-293 cells were engineered in our laboratory using CRISPR-Cas technology for the deletion of all of the three-native endogenous IP_3_R sub-types (HEK-3KO) (31). HEK-3KO cells were cultured in Dulbecco’s modified Eagle’s medium supplemented with 10% fetal bovine serum, 100 U/mL penicillin, and 100 μg/mL streptomycin in an incubator set to 37°C with 5% CO_2_. Transfection of WT IP_3_R1, E1128A or E1131A plasmid construct was performed using Lipofectamine 2000 (#11668019, Invitrogen). Cells were allowed to recover for 48 h, and subsequently sub-cultured in new 60 mm plates containing media supplemented with 1 mg/mL G418. Following 7 days of selection, individual colonies of cells were picked and transferred to new 24-well plates containing media supplemented with 1 mg/mL G418. Clonal lines were expanded, and the expressing of desired constructs were confirmed by western blotting.

### Western blotting

Total protein was isolated from indicated HEK/DT40 control and stable cell lines using membrane-bound extraction buffer (10 mM Tris_HCl, 10 mM NaCl, 1 mM ethylene glycol tetra acetic acid (EGTA), 1 mM ethylenediaminetetraacetate, 1 mM NaF, 20 mM Na_4_P_2_O_7_, 2 mM Na_3_VO_4_, 1% Triton X-100 (vol/vol), 0.5% sodium deoxycholate [wt/vol], and 10% glycerol) supplemented with protease inhibitors (#78429, Thermo Scientific). For protein isolation, following the addition of an appropriate amount of lysis buffer, cells were harvested in 1.5 mL tubes and placed on ice for 30 min. To disrupt the cells, the tubes were vortexed for 10 s every 10 min and returned on ice. Following incubation on ice, the cell lysates were centrifuged at 13,000 rpm at 4°C for 10 min. The supernatant was transferred to new labeled tubes. Protein concentration in the lysates was estimated using the Dc protein assay kit (Bio-Rad). Equal amounts of lysates along with a protein standard marker (#161-0374, Biorad) were then subjected to sodium dodecyl sulfate-polyacrylamide gel electrophoresis (SDS-PAGE) and transferred to a nitrocellulose membrane. After blocking, the membranes were incubated with the indicated primary antibodies and appropriate secondary antibodies before imaging with an Odyssey infrared imaging system (LICOR Biosciences). Band intensities were quantified using Image Studio Lite version 5.2 and presented as ratios of IP_3_R1 to glyceraldehyde 3-phosphate dehydrogenase (GAPDH). IP_3_R1 antibody was generated by Antibody Research Corporation and used at 1:1000 dilution; GAPDH (#AM4300, 1:75,000 dilution), secondary goat anti-rabbit (SA535571) and secondary goat anti-mouse (SA535521) antibodies were from Invitrogen and used at 1:10,000 dilution (14).

### Immunocytochemistry and confocal microscopy

HEK-3KO and HEK-3KO cells stably expressing the WT and mutant E1128A, E1131A IP_3_R1 constructs were plated on poly-D-lysine (100 μg/mL)-coated coverslips. At roughly 70% confluent, cells were fixed using 4% paraformaldehyde at room temperature for 10 min. Subsequently, the coverslips were washed with PBS, and cells were blocked in 10% bovine serum albumin (BSA) for 1 h at room temperature. Following blocking, cells were incubated overnight at 4°C with primary antibody against IP_3_R1 (#ARC154, Antibody Research Corporation, 1:1000 dilution). The following day, the primary antibody was removed, and coverslips were washed 3 times with PBS for 10 min with gentle rocking. Subsequently, the cells were incubated with Alexa Fluor 488 goat anti-rabbit IgG secondary antibody (#A11008, Invitrogen, 1:1000 dilution) was incubated for 1 h at room temperature with gentle rocking. After incubation, coverslips were washed with PBS and mounted on slides. After allowing the slides to dry, coverslips were sealed onto slides and imaged using confocal microscopy using an Olympus Fluoview 1000 microscope (27).

### Measurement of cytosolic Ca^2+^ in intact cells

Population-based Ca^2+^ imaging in HEK-3KO and HEK-3KO cells stably expressing the WT and mutant E1128A, E1131A IP_3_R1 constructs was performed as described previously (52). Briefly, adherent cells were cultured in 10 cm^2^ cell culture dishes. Upon attaining 90 to 100% confluency, the cells were loaded with 4 μM Fura-2/AM in cell culture media and incubated at 37°C in the dark for 1 h. The cells were subsequently washed 3 times with imaging buffer (10 mM HEPES, 1.26 mM Ca^2+^, 137 mM NaCl, 4.7 mM KCl, 5.5 mM glucose, 1 mM Na_2_HPO_4_, and 0.56 mM MgCl_2_, at pH 7.4). An equal number (200,000 cells/well) of cells were seeded into each well of a black-walled 96-well plate. Fura-2/AM imaging was carried out by alternatively exciting the loaded cells between 340 and 380 nm; emission was monitored at 510 nm using FlexStation 3 (Molecular Devices). Peak response to various concentrations of CCh (0.1–100 μM) was determined using SoftMax Pro Microplate Data Acquisition and Analysis software as described previously (52). Data from at least three individual plates on different days after curve fitting using a logistic dose-response equation in GraphPad Prism 10 are presented.

Single-cell Ca^2+^ imaging in the aforementioned cells was performed as described previously (52). Briefly, cells were seeded on 15-mm glass coverslips in 12-well plates and left undisturbed for 24 hours. Once attached to the coverslip, the cells were washed with imaging buffer before attachment of the glass coverslip to a Warner perfusion chamber using vacuum grease. Subsequently, the cells were loaded with 2 μM Fura-2/AM for 25 min in the dark at room temperature. Cells were then perfused with Ca^2+^ containing imaging buffer and stimulated with indicated concentrations of CCh in Ca^2+^ containing or Ca^2+^ free imaging buffer (10 mM HEPES, 1 mM EGTA, 137 mM NaCl, 4.7 mM KCl, 5.5 mM glucose, 1 mM Na_2_HPO_4_, and 0.56 mM MgCl_2_, at pH 7.4). For single-cell Ca^2+^ imaging using DT40 stable cells, pelleted cells were loaded with 2 μM Fura-2/AM and either 1 or 5 μM ci-IP_3_/PM (Tocris #6210) on 15-mm glass coverslips for an hour at room temperature. To photo-release IP_3_, ultraviolet (UV) light from Photonics UV-Flash II Pulsed Light Source was introduced to uniformly illuminate the field of view. Ca^2+^ imaging was performed using an inverted epifluorescence Nikon microscope equipped with a 40× oil immersion objective. Fura-2/AM imaging was carried out by alternatively exciting the loaded cells between 340 nm and 380 nm; emission was monitored at 505 nm. Images were captured every second with an exposure of 20 ms and 4 × 4 binning using a digital camera driven by TILL Photonics software as previously described (52). Image acquisition was performed using TILLvisION software and data were exported to Microsoft Excel, where data were analyzed for change in peak amplitude, percentage of cells with predefined peak amplitudes (340/380>0.1), and fraction of oscillating cells (Δ340/380>0.05). Each experiment was repeated at least three times on different days.

### Detection and analysis of Ca^2+^ puffs using TIRFM

HEK-3KO cells stably expressing the WT and mutant E1128A, E1131A IP_3_R1 constructs were cultured on 15-mm glass coverslips coated with poly-D-lysine (100 μg/mL) in a 35-mm dish for 36 h. The cells were washed three times with imaging buffer. Subsequently, the cells were incubated with 5 μM Cal520-AM (#21130, AAT Bioquest) and 0.05 μM ci-IP_3_/PM (Tocris #6210) in imaging buffer supplemented with 0.01% BSA in dark at room temperature. After 1 h incubation, the cells were washed 3 times with imaging buffer and incubated in imaging buffer containing 5 μM EGTA-AM (#E1219, Invitrogen). After 45 min of incubation, the media was replaced with fresh imaging buffer and incubated for an additional 30 min at room temperature to facilitate de-esterification of loaded reagents.

Next, the coverslip was mounted in a warner chamber and imaged using an Olympus IX81 inverted TIRFM equipped with oil-immersion PLAPO OTIRFM 60× objective lens/1.45 numerical aperture. Olympus CellSens Dimensions 2.3 (Build 189987) software was used for imaging. The cells were illuminated using a 488-nm laser to excite Cal-520 and the emitted fluorescence was collected through a band-pass filter by a Hamamatsu ORCA-Fusion complementary metal oxide semiconductor camera. The angle of the excitation beam was adjusted to achieve TIRF with a penetration depth of ∼140 nm. Images were captured from a final field of 86.7 × 86.7 μm (400 × 400 pixels, 1 pixel = 216 nm) at a rate of ∼50 frames/second (binning 2 × 2) by directly streaming into random access memory. To photo-release IP_3_, ultraviolet (UV) light from a laser was introduced to uniformly illuminate the field of view. Images were exported as .vsi files. Images, 5 s before and 60 s after flash photolysis of ci-IP_3_, were captured, as described previously (53). The .vsi files were converted to .tif files using Fiji and further processed using FLIKA, a Python programming-based tool for image processing (54). From each recording, ∼100 frames (∼2 s) before photolysis of ci-IP_3_ were averaged to obtain a ratio image stack (F/F_o_) and standard definition for each pixel for recording up to 13 s following photolysis. The image stack was Gaussian filtered, and pixels that exceeded a critical value (1.0 for our analysis) were located. The “Detect-puffs” plug-in was used to detect the number of clusters (puff sites), number of events (number of puffs), and the amplitudes and durations of localized Ca^2+^ signals from individual cells. All of the puffs identified automatically by the algorithm were manually confirmed before analysis. The results from FLIKA were saved to Microsoft Excel and graphs were plotted using GraphPad Prism 10 (54).

### Preparation of DT40 Cell Nuclei

Isolated DT40 nuclei were prepared using homogenization. The homogenization buffer (HB) contained 250 mM sucrose, 150 mM KCl, 3 mM 2-mercaptoethanol (β-ME), 10 mM Tris, 1 mM phenylmethanesulphonylfluoride, pH 7.5, with a complete protease inhibitor tablet (Roche). Cells were washed and resuspended in HB before nuclear isolation using an RZR 2021 homogenizer (Heidolph Instruments) with 15 strokes at 1,200 rpm. A 3-μL aliquot of nuclear suspension was placed in a 3-mL bath solution, which contained 140 mM KCl, 10 mM HEPES, 500 μM BAPTA (1,2-bis(o-aminophenoxy)ethane-N,N,N0,N0-tetraacetic acid), and 246 nM free Ca^2+^, pH 7.1. Nuclei were allowed to adhere to a plastic culture dish for 10 min before patching (27).

### On-Nuclei Patch-Clamp Experiments

Single IP_3_R1 channel potassium currents (i_k_) were measured in the on-nucleus patch-clamp configuration using pCLAMP 9 and an Axopatch 200B amplifier (Molecular Devices). Pipette solution contained 140 mM KCl, 10 mM HEPES, either 5 mM or 100 µM Na-ATP, with varying concentrations of IP_3_ (1 µM or 10 µM), BAPTA, and free Ca^2+^. Free [Ca^2+^] was calculated using Max Chelator freeware and verified fluorometrically. Traces were consecutive 3-s sweeps recorded at −100 mV, sampled at 20 kHz, and filtered at 5 kHz. A minimum of 15 s of recordings were considered for data analyses. The data are representative of between three and five experiments for each condition presented. Pipette resistances were typically 20 MΩ and seal resistances were >5 GΩ (27).

### Data Analysis

Single-channel openings were detected by half-threshold crossing criteria using the event detection protocol in Clampfit 9. We assumed that the number of channels in any particular nuclear patch is represented by the maximum number of discrete stacked events observed during the experiment. Even at low Po, stacking events were evident. Only patches with one apparent channel were considered for analyses. Probability of opening (Po), unitary current (ik), open and closed times, and burst analyses were calculated using Clampfit 9 and Origin 6 software (OriginLab). All-points current amplitude histograms were generated from the current records and fitted with a normal Gaussian probability distribution function. The coefficient of determination (R2) for every fit was >0.95. The Po was calculated using the multimodal distribution for the open and closed current levels. The threshold for an open event was set at 50% of the maximum open current and events shorter than 0.1 ms were ignored. A “burst” was defined as a period of channel opening following a period of no channel activity, which was >5 times the mean closed time (0.2 ms) within a burst. Ca^2+^ dependency curves were fitted separately for activation and inhibition with the logistic equation:

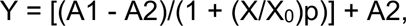

where A1 and A2 are asymptotes, X is the concentration of Ca^2+^, X_0_ is the half-maximal concentration, and p is the slope related to the Hill coefficient. Equation parameters were estimated using a nonlinear, least-squares algorithm.

### Statistical Analysis

All of the statistical tests were conducted in GraphPad Prism 10 and data are presented as the mean±s.e.m. Statistical significance was determined as indicated in the figure legends.

## Acknowledgements

The authors wish to thank all the members of Yule lab for their valuable suggestions.

